# DAISIEprep: an R package for the extraction and formatting of data for the island biogeography model DAISIE

**DOI:** 10.1101/2023.02.19.529129

**Authors:** Joshua W. Lambert, Lizzie Roeble, Théo Pannetier, Rampal S. Etienne, Luis Valente

**Author notes:** Corresponding Author: Joshua W. Lambert, Groningen Institute for Evolutionary Life Sciences, Box 11103, 9700 CC Groningen, The Netherlands. These authors contributed jointly to senior authorship (Rampal S. Etienne, Luis Valente).

## Abstract

1. Phylogenetic trees are commonly used to answer questions on biogeographical and diversification histories of different groups.
2. Recently, new approaches have been developed that use community phylogenetic trees requiring a data structure distinct from the single phylogenetic trees that are commonly used, which may be a barrier to the utilisation of these approaches.
3. DAISIE (Dynamic Assembly of Islands through Speciation, Immigration and Extinction) is an island biogeography model that can estimate rates of colonisation, speciation and extinction from phylogenetic data across insular communities, as well as simulate islands under those rates.
4. Here we describe the `DAISIEprep` R package, a set of pre-processing tools to aid the extraction of data from one or many phylogenetic trees to generate data in a format interpretable by DAISIE for the application of island biogeography inference models. We present examples to illustrate the various data types that can be used.
5. The package includes simple algorithms to extract data on island colonists and account for bio-geographical, topological and taxonomic uncertainty. It also allows flexible incorporation of either missing species or entire insular lineages when phylogenetic data are not available.
6. `DAISIEprep` enables reproducible and user-friendly data extraction and formatting, and will facilitate addressing questions about island biogeography, diversification and anthropogenic impacts in insular systems.

## Introduction

Phylogenetic trees are prevalent in evolutionary, epidemiological, and even linguistic sciences, and typically use well-defined data structures. Standard tree data structures, particularly the commonly used ‘Newick’ format (Maddison *et al*. 1997; Felsenstein 2004; Cardona *et al*. 2008), are employed within and across programming languages. For example, in the R language, phylogenetic tree formats have been constructed that are compatible with different packages and methodologies (Paradis *et al*. 2004; FitzJohn 2012; Revell 2012; Pennell *et al*. 2014; Morlon *et al*. 2016; R Core Team 2022). However, newly developed methods that use phylogenetic information may require different data structures, potentially creating obstacles for the scientific community to apply those methods. In particular, island biogeography research is increasingly utilising phylogenetic information to answer questions on island community assembly and diversification dynamics, but it requires phylogenetic information in an unconventional format. Instead of relying on a single phylogenetic tree representing a clade of interest, it can comprise data assembled from multiple phylogenetic trees of different clades and even non-phylogenetic information. In contrast to the phylogenetic tree formats, standards and methods for manipulating island community data are lacking.

DAISIE (Dynamic Assembly of Island biota through Speciation, Immigration and Extinction) is a model of island biogeography and macroevolution that focuses on entire island communities or assemblages, complementing a growing body of work that integrates multiple distinct clades within a given community (Webb *et al*. 2002; Cavender-Bares *et al*. 2009; Silvestro *et al*. 2015; Condamine *et al*. 2019). The model uses times of island colonisation and speciation events as well as the endemicity status of the island species to estimate a set of parameters (cladogenesis rate *λ*^*c*^, extinction rate *µ*, colonisation rate *γ*, anagenesis rate *λ*^*a*^ and a carrying capacity *K*^*′*^). The framework is founded on the birth-death model for reconstructed phylogenetic trees (i.e. only extant species) and can incorporate diversity-dependence in colonisation and speciation rates through a clade-specific (only within clades) or island-wide (across the entire island community) carrying capacity, as well as species known to exist but not present in the phylogeny (Etienne *et al*. 2012; 2022). DAISIE has been used for various studies in island biogeography. For example, it has allowed estimating rates of speciation, colonisation and extinction of the Galapágos terrestrial avifauna (Valente *et al*. 2015), it enabled the first global analysis of MacArthur & Wilson’s (1967) area and isolation model using phylogenetic data (Valente *et al*. 2020), it has put the impact of human-induced extinction in a macro-evolutionary context (Michielsen *et al*. 2023), and it has shown the presence of colonisation rate shifts of fishes in a Japanese Lake (Hauffe *et al*. 2020).

DAISIE allows studying diversification dynamics in insular communities without the need for a full phylogeny. Such a phylogeny is often not available, and the often employed alternative, focusing on the phylogeny of a single clade on the island, will not make use of the information from other lineages (small or large), and cannot be used to estimate colonisation rates. DAISIE uses a unique data structure: instead of considering a single insular lineage (e.g., the phylogeny of the Galápagos finches), it focuses on multiple independent lineages descending from different colonisation events of the island, representing various lineages that make up an insular community (e.g. phylogenies for each of the birds that colonised the Galápagos). Some of these lineages may have radiated into a large clade with many species on the island, others may have only a single species or few species on the island. Some of the species may be endemic to the island (found only on the island) and some may be non-endemic (also present outside of the island, i.e. the mainland). Phylogenetic data of sister species on the mainland are also needed to obtain an estimate of the colonisation time for each lineage. The data used by DAISIE thus includes colonisation and speciation times for the various lineages (obtained from time-calibrated molecular phylogenies), endemicity status (endemic or non-endemic to the island) and the number of missing species (how many species from each insular lineage are not represented because molecular data is not available). A key feature of the model is that it does not require all the information on the different insular lineages to come from a single phylogeny. Instead, data from different phylogenetic trees can be used (for example, phylogenetic analyses of different genera or families from different publications).

The specific format of the data required to fit DAISIE models (Figure 1B) may constitute a barrier to its use. Here we present an R package, `DAISIEprep`, to provide a comprehensive set of tools to easily create data objects from phylogenetic data and island species checklists to be analysed in the maximum likelihood island biogeography model DAISIE (Lambert *et al*. 2023b). The package facilitates reproducible, fast extraction of community data from phylogenetic data (Figure 1). It is also possible to insert data on species or clades for which phylogenetic data is not available, but for which the number of species and endemicity statuses are known. If a species or island lineage is not sampled in a time-calibrated phylogeny, but an estimate for the time of island colonisation is available from other data sources, this can also be included. As software that produces phylogenetic data – for example BEAST2 (Bouckaert *et al*. 2014) and MrBayes (Huelsenbeck & Ronquist 2001) among others – output standard phylogenetic data structures, a package to convert these types of data into island-specific community data is a useful intermediate tool between primary empirical data and model fitting. Here, we explain the main functionalities of the R package across a range of scenarios, some auxiliary functions, as well as the downstream influence of data extraction choices.

**Figure 1:**
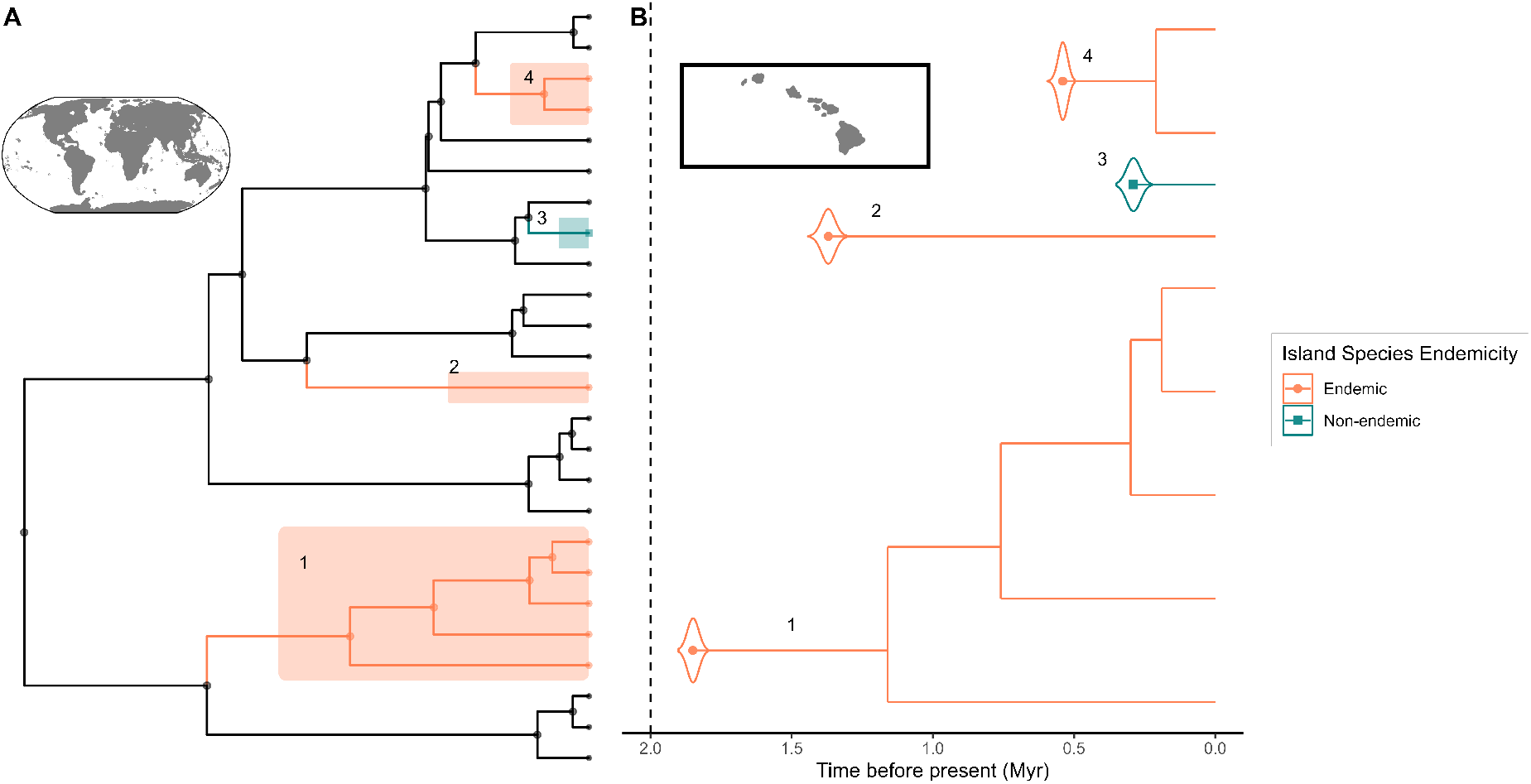
Visual representation of a typical dataset used in DAISIE analyses. In this hypothetical example, the focal insular system is the archipelago of Hawaii, and the focal island community has nine species. Panel A shows the wider global phylogeny in which the nine Hawaiian species are embedded (highlighted). Panel B shows the separate phylogenies of those island species from the perspective of the archipelago. Numbers on the plots link the corresponding lineages from each panel. The island community (B) consists of lineages resulting from four colonisation events, which are spread out in different topological locations in the tree of the taxonomic group at the global scale (A). Three of the island lineages contain exclusively species that are endemic to the island and one contains only a single non-endemic species. Two of the endemic lineages have undergone cladogenetic speciation on the island, forming island clades with more than one species. The colonisation time is assumed to be the divergence time from the mainland sister species (stem age). Violin plots around the island colonisation time show the uncertainty in divergence time estimates, representing a range of times from a posterior distribution of trees. The objective of the `DAISIEprep` package is to convert global phylogenetic data (A) to island community data (B).

## Package description

The package aims to facilitate and automate extracting and formatting data prior to applying DAISIE. This alleviates users having to write custom scripts. The `DAISIEprep` package requires as input a checklist of species on the island or archipelago of interest as well as time-calibrated phylogenetic data for these species. `DAISIEprep` uses phylogenetic data stored as a `phylo4d` object from the `phylobase` R package (Hackathon *et al*. 2020) which can handle phylogenetic trees with data at the tips and nodes of the tree. We also introduce three novel data structures (`Island_colonist`, `Island_tbl` and `Multi island tbl`) to aid in the handling of the specific community phylogenetic data, as well as a set_of_functions (methods) to easily modify the data, making use of the benefits of object-oriented programming in R (see Chambers (2014)). To check if the input data is compatible with `DAISIEprep`, the function `check_phylo_data()` can be used.

The package provides two algorithms to extract the community phylogenetic data: (1) minimum time of colonisation (‘min’), and (2) ancestral state reconstruction (‘asr’). The ‘min’ algorithm assumes that the stem node of an island lineage (the split from closest non-island or island non-endemic relatives sampled in the phylogeny) corresponds to the colonisation time of the island; and that there is no back-colonisation from the island to the mainland. The ‘min’ algorithm’s assumption that species cannot migrate away from the island are consistent with the assumptions of the original DAISIE model. Under these assumptions, colonisation times are taken as the stem age. In the case when non-endemic island species or endemic island species with no sister species on the island are represented by a single tip in the phylogeny, the age of the node leading to the single tip corresponds to the colonisation time. When they are represented by several individuals or tips in the phylogeny, and in the case of endemic island clades with more than one species sampled in the phylogeny, the stem age of the island lineage corresponds to the colonisation time. This algorithm is useful when data seems to abide by the assumption of no back-colonisation or colonisation to other islands thus changing the endemicity status to non-endemic. However, should such colonisation occur, the estimated number of island colonisations will be inflated, as island radiations may be artificially broken up into multiple colonisations of the island. In such cases, one may prefer to extract species based on a different model. To determine when colonisation happened, this would require a probabilistic mapping of island occupancy along the internal branches of the phylogeny, i.e. ancestral state reconstruction (see Joy *et al*. (2016) for review). The ‘asr’ algorithm is based on phylogenetic ancestral state reconstruction, and uses the ancestral geographical states at the nodes within the tree inferred with reconstruction methods to determine a likely number of colonisation events of the island. `DAISIEprep` provides the `add_asr_node_states()` function to easily estimate ancestral states, using the `castor` R package to provide the maximum parsimony and the Mk model (Louca & Doebeli 2018).

The `DAISIEprep` package is set up to allow users to easily import ancestral range reconstructions obtained using methods from other packages, such as DEC/DEC+J (Matzke 2014; 2021), State-Speciation-Extinction models (Maddison *et al*. 2007; Goldberg *et al*. 2011; FitzJohn 2012; Herrera-Alsina *et al*. 2019), or stochastic character mapping (Huelsenbeck *et al*. 2003). More details are provided in the ancestral state reconstruction extensions vignette in the `DAISIEprep` package.

The ‘asr’ algorithm prevents clades being artificially split up in the case of a further colonisation event (Figure 2). In the extraction process the non-endemic is represented as an endemic species in the `Island_tbl` and when fitting the DAISIE model. Therefore, the differences between the ‘min’ and ‘asr’ algorithms are both in the number of colonisation events inferred, as well as the endemicity of species within island clades (Figure 2). Furthermore, under both ‘min’ and ‘asr’ algorithms, it is possible that the phylogenetic data or state reconstruction indicate a colonisation time older than the island age, for example if the closest relative of the island species has gone extinct or is not sampled in the phylogeny (Lambert *et al*. 2022). In such cases, the island lineage is assumed to have colonised any time after the island formation, with the DAISIE inference model integrating over this uncertainty.

**Figure 2:**
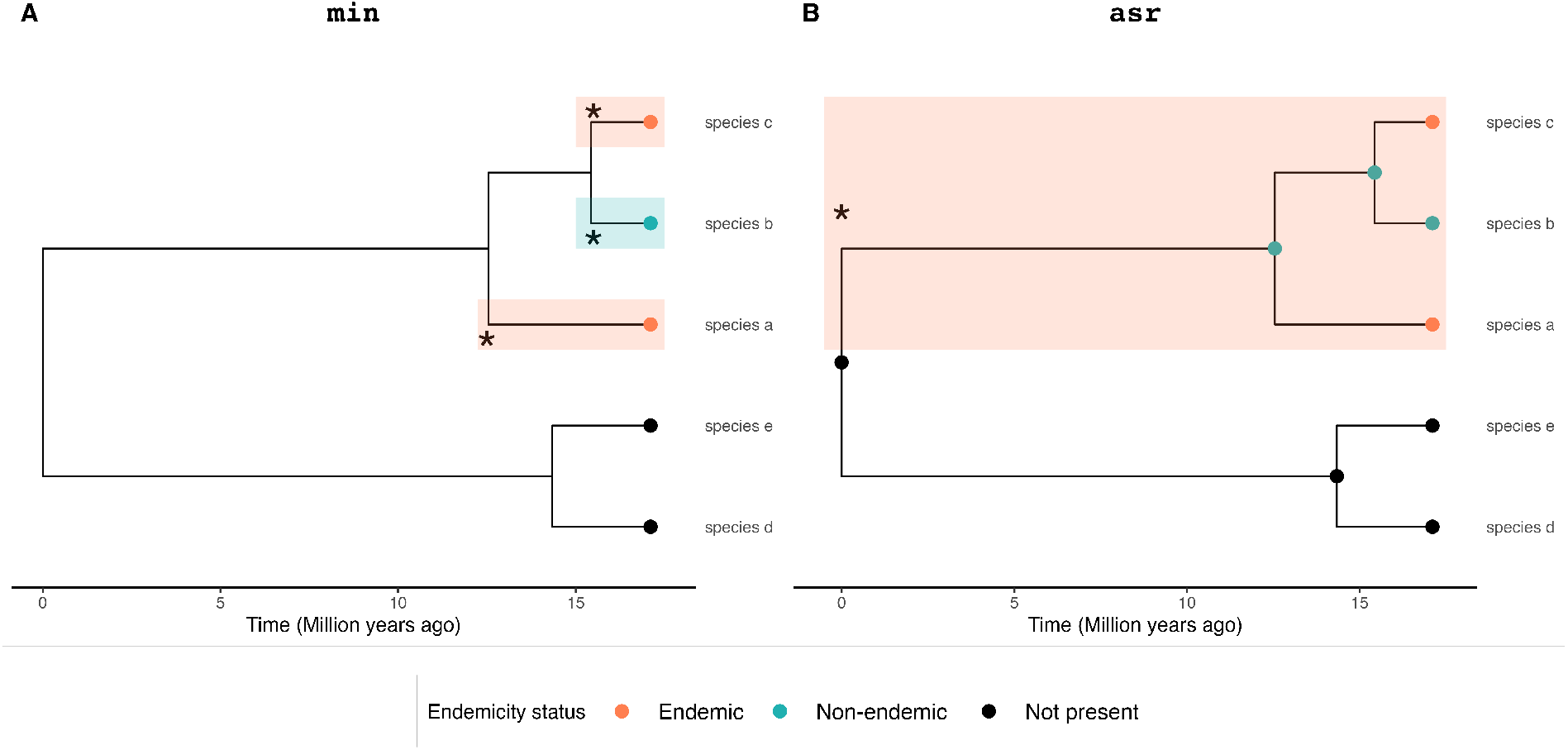
Comparison of the ‘min’ (A) and ‘asr’ (B) algorithms for data extraction. Two identical phylogenetic trees with five species, two of which are endemic, one non-endemic and two not present on the island. The ‘min’ algorithm (A) extracts three species, with stem times shown by **\*** and shaded areas, as there is a non-endemic species embedded within a clade with two endemic species. Thus using the ‘min’ algorithm assumes independent colonisations of each, minimising the time of colonisation. In contrast, the ‘asr’ algorithm (B) uses the internal node states (shown by dots at each node), to extract a single island colonist with three species, with the stem age of the island clade assumed to be the colonisation time (**\*** and shaded area). In this case (B), the extracted clade (shaded) is assumed endemic even though it contains a non-endemic species given the assumptions of the DAISIE inference model. Therefore the endemicity status of the non-endemic is converted to endemic because the DAISIE inference model does not allow non-endemic species embedded within endemic island clades from a single colonisation because it does not consider colonisation from the island.

### Installation of DAISIEprep

The `DAISIEprep` package can be installed from github using the `remotes` R package (Csárdi *et al*. 2021) `remotes::install_github(“ joshwlambert/DAISIEprep”`) and will be available on CRAN.

## Applications

An analysis using the DAISIE model starts with a checklist of the (native) species of the focal group that are found on an island, for example, all ferns or all squamates. This checklist should include whether each species is endemic to the island or not. The next step is to gather phylogenetic data from which colonisation and branching times of the island species can be inferred. The ‘ideal’ dataset would be a single time-calibrated phylogenetic tree with complete sampling, in which all species of the focal group are included (preferably multiple individuals of the same species) as well as all of their closest non-island relatives. In practice this type of dataset is very rare and difficult to produce. For most insular communities, there may be a few well-sampled phylogenies available for some of the insular lineages, while other lineages may have poorly sampled phylogenies, and for some lineages there may be no dated phylogenetic data available at all. For some non-endemic species there may be individuals from the mainland sampled in the phylogeny, but not from the island. DAISIE can incorporate information from these heterogeneous data types.

Here we demonstrate the basic usage of the `DAISIEprep` package through two applications to empirical data. The first is on a species-level macro-phylogeny of mammals by Upham *et al*. (2019), which we use to obtain information on the mammals of the island of Madagascar. This represents an application that may be common when users want to extract island data from a single global/macro-phylogeny containing a whole taxonomic group (e.g. birds (Jetz *et al*. 2012), or amphibians (Jetz & Pyron 2018)). The second empirical example is a data set of the Asteraceae plant family of the Hawaiian archipelago. Asteraceae form a hyper-diverse group of plants that is globally distributed on islands. Unlike for mammals, there is no species-level phylogenetic tree for the Asteraceae family (*>* 30, 000 species). Thus, the Hawaiian Asteraceae dataset consists of multiple phylogenies for different Hawaiian genera or radiations, as well as species richness and endemism data for those Hawaiian clades for which no phylogenetic data are available (Landis *et al*. 2018; Knope *et al*. 2020; Keeley *et al*. 2021). Together, these two examples illustrate the use of macro-phylogenies, single or multiple phylogenies as well as the addition of island taxa when no phylogenetic data is available.

### Single phylogeny example: mammals of Madagascar

First, we used the checklist of all the mammal species that inhabit Madagascar from Michielsen *et al*. (2023), which recorded the island species’ endemicity status (endemic or not endemic to Madagascar). A table with this endemicity information is a requirement for all uses of the `DAISIEprep` package. We link this table with the phylogenetic data using the `create_endemicity_status()` function and the abovementioned `phylo4d` class (this creates the `phylod` object in the code example below). The mammal phylogeny used is the maximum clade credibility tree (Heled & Bouckaert 2013) from Upham *et al*. (2019). For this example we will use the ‘asr’ algorithm. The table (checklist of species with endemicity statuses) and phylogeny are all the data required to extract the island community data using `extract_island_species()`.

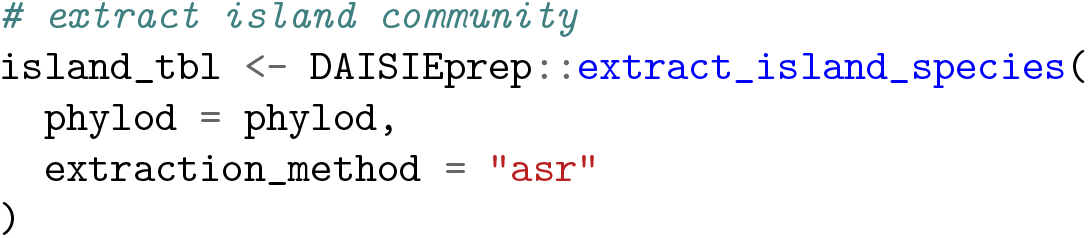

This outputs an `Island_tbl` object containing all the information on the colonisation and island speciation times, the endemicity status, the number of missing species and name of each colonist. By default, the number of missing species for each island colonist is set to zero. To automatically count and assign missing species to the `Island_tbl` using information stored in the island species checklist, `count_missing_species()`, `unique_island_genera()` and `add_multi_missing_species()` can be used. Conversely, if missing species have been misassigned in this process, they can manually be removed from the `Island_tbl` using `rm_multi_missing_species()`. This set of functions assists in tabulating and assigning missing species to the island community data that was directly extracted from the phylogenetic tree (using `extract_island_species()`). To deal with the missing species remaining after automatic assignment, we provide a set of functions to add the missing species manually, in one of two ways, either assigning the missing species to an existing island clade (not to a specific topological location within the clade, but contributing to the total diversity of the clade) in the `Island_tbl` using `add_missing species()`, or adding the missing species as a new separate island clade using `add island_colonist()` (Figure 3). See the tutorial vignette in the `DAISIEprep` R package for worked examples of adding missing species under the different scenarios.

**Figure 3:**
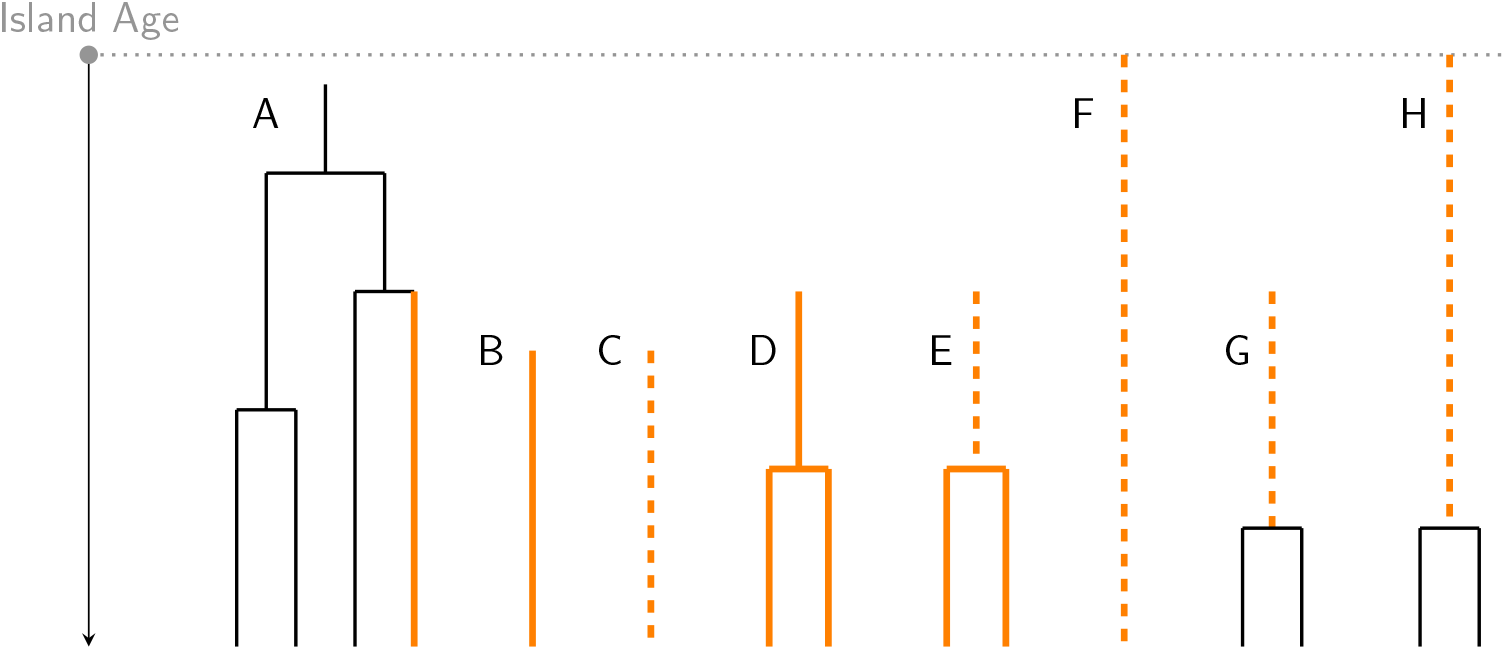
Adding missing species to island community data. The clade labelled A is a case when the phylogeny of an island clade is known (black lines), but one of the species is missing from the phylogeny (orange line). In this scenario the `add_missing_species()` function would be used to add, in this case, one missing species to the existing island clade in the island data set. The lineage labelled B is an island singleton (i.e. not an island radiation/clade) which is known to be on the island but has no time-calibrated phylogenetic data available. In this case an estimate of time of island colonisation exists, possibly from the literature. The lineage labelled C is the same as B except that the time taken from the literature is considered a maximum possible colonisation time rather than an exact time of colonisation (indicated by the dashed line). Island clade D is the same as B except that there is a clade instead of a singleton, in this case two species, on the island. Island clade E is the same as D but the colonisation time is a maximum and not an exact time (as in C). Lineage F is the case when a species is known on the island but its time of colonisation is unknown. Island clades G and H represent cases where the time of colonisation from the stem age is known or unknown, respectively, for a clade of two island species but an island speciation time is known (crown age) and can be used as a minimum time of colonisation. When the DAISIE model is fitted to the cases shown in C, E, F and H the model integrates between the maximum and minimum possible colonisation times (dashed lines) Examples B-H all use the `add_island_colonist()` to add island lineages the data set. The topology of missing species (island clade A) and their branching times (island clades D and E) are not known and are shown here just for illustrative purposes; see Etienne *et al*. (2012). Species endemicity is not shown for simplicity. See the tutorial vignette in the R package for a full explanation of each case with examples.

When using `add_island_colonist()`, a colonisation can either be taken from the literature, extracted from the stem age of the genus if it is present in the phylogeny (i.e. the genus has mainland species in the tree) (Figure 3B, C, D, E), or can be given as unknown (using `NA`) (Figure 3F). An example of adding missing species in the case of the Malagasy mammals is the bat genus *Chaerephon*, which is missing and needs to be assigned, but it has species in the phylogeny not present on Madagascar. Thus a stem age can be extracted which is used as a maximum possible colonisation time. Setting `col_max_age = TRUE` specifies that the colonisation time given (in this case, the stem age) should be considered an upper bound rather than a precise time of colonisation.

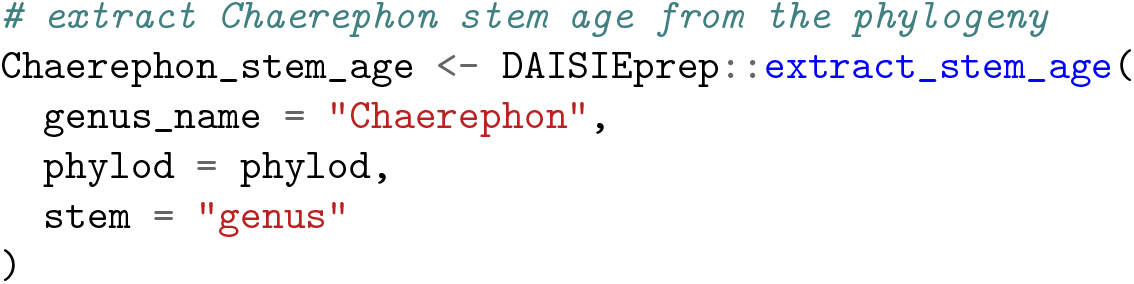

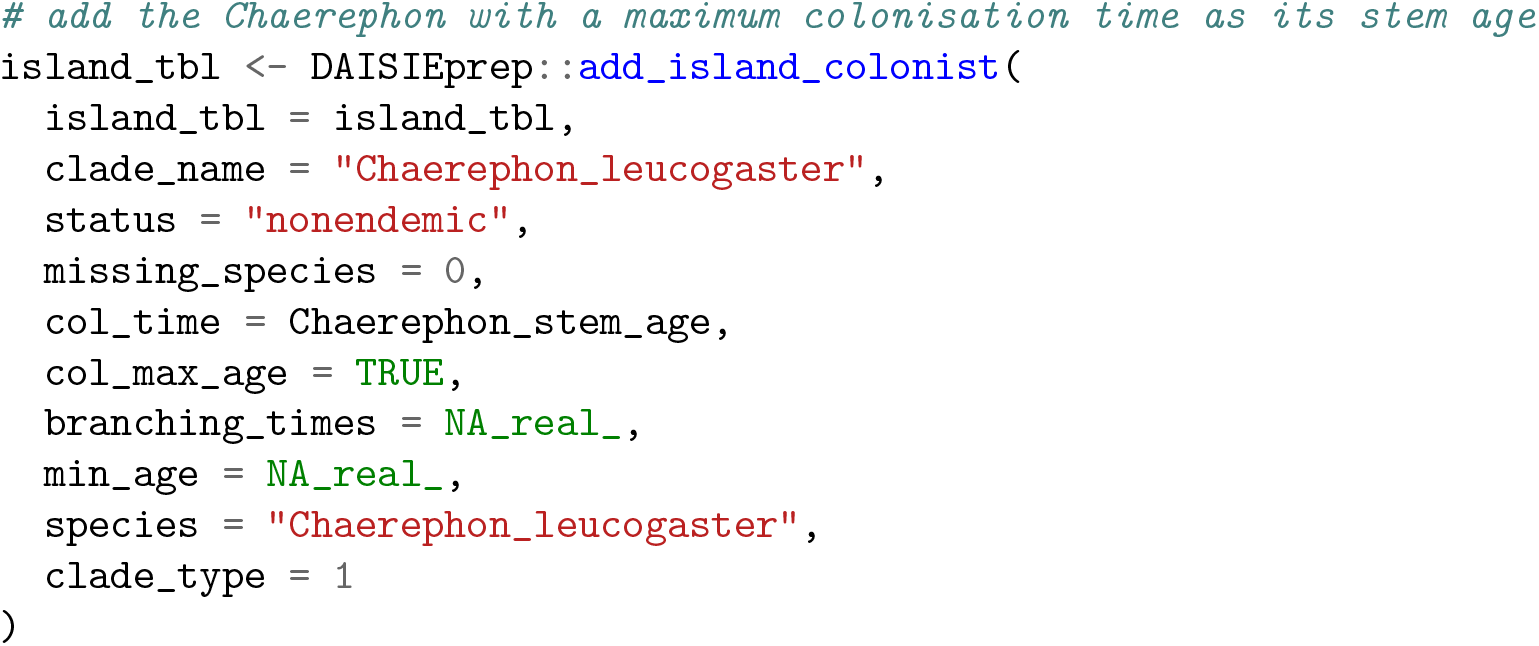

Once the `Island_tbl` object is finalised it can be converted to the DAISIE data list structure required by the DAISIE likelihood model, using `create_daisie_data()` function. The script and phylogenetic data used for the mammals of Madagascar in this section for loading the checklist, extracting the data, and handling the missing species can be found in the `DAISIEprepExtra` R package (Lambert *et al*. 2023a).

### Multiple phylogenies example: Hawaiian Asteraceae

The case of the Asteraceae species of the archipelago of Hawaii provides a good example of the spectrum of phylogenetic and diversity data available for an island community. The Hawaiian archipelago currently has 98 native species (97 endemic) of Asteraceae, which are the result of at least 10 colonization events. There are several new species that are currently awaiting description, so this number will change in the future, but for the purposes of this example we use the currently accepted species. Each of the Hawaiian Asteraceae clades has received varying degrees of attention, contributing to differing data availability across the community. For example, the Hawaiian Silverswords and Pacific *Bidens* – two classic examples of adaptive radiations – are well-studied with dated phylogenies available; whereas, *Tetramolopium* – a potential radiation of 11 species – does not have a phylogeny available.

We compiled a checklist of the Asteraceae species native and endemic to Hawaii and gathered all available phylogenetic data. For the 10 assumed colonization events, three clades (representing 60.5% of the species) have dated phylogenies publicly available (Hawaiian Silversword alliance, Landis *et al*. (2018); *Bidens*, Knope *et al*. (2020); *Hesperomannia*, Keeley *et al*. (2021)); three clades have dated phylogenies but tree files are not publicly available, so we use the colonisation time estimates cited in the text of the publications (*Artemisia*, Hobbs & Baldwin (2013); *Lipochaeta-Melanthera* alliance, Edwards *et al*. (2018); *Pseudognaphalium*, Nie *et al*. (2016)); and four clades do not have a published phylogeny, and we use the diversity data (number of species thought to belong to the same insular lineage based on taxonomic information, and endemicity status) (*Tetramolopium, Keysseria, Remya, Adenostemma*). The resulting community dataset leverages multiple dated phylogenies, colonisation time estimates provided in-text in scientific articles, and island diversity data (number of species and endemicity status) for cases for which no phylogenetic work has been done on the island species or clades (Figure 4). For the cases where only crown ages were available, these were used as the minimum colonisation times (most recent possible time of colonisation).

**Figure 4:**
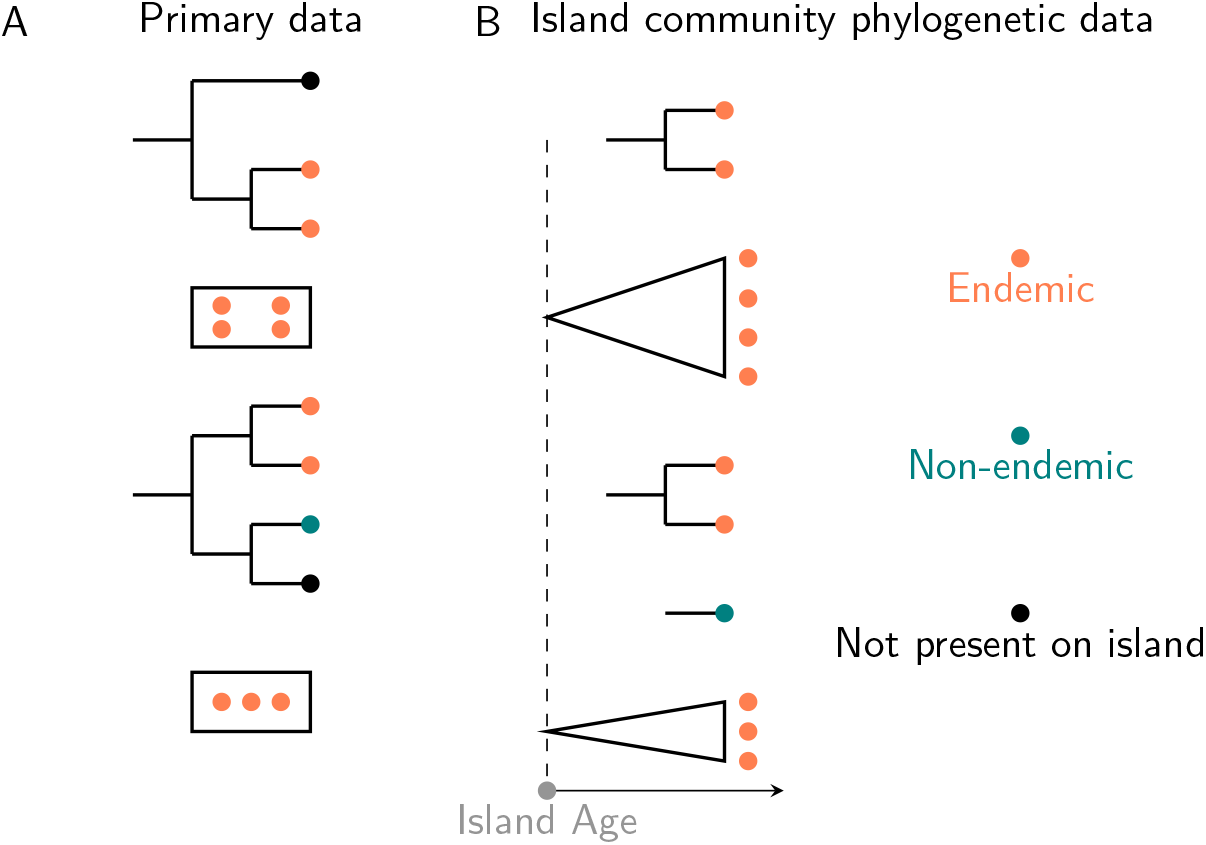
Example of using different sources and formats of data to obtain island community phylogeny data. In this case, the primary data (A) is composed of two phylogenetic trees that include some of the island species and, for cases where phylogenetic data is not available (represented by rectangular boxes), diversity data (number of species present on the island assumed to belong to the same insular lineage, and their endemicity status). This example shows how island species or clades that are missing phylogenetic data but for which the island species diversity and endemism are known (shown in boxes), can be accounted for in the DAISIE framework (B) by considering them as missing species and treating the clade as a polytomy with known species richness and species endemism (shown by the triangles with the dots representing how many species are in those clades). The dashed line represents the age of the island, and those island species that lack phylogenetic data are assumed to have colonised the island any time after island formation.

The pipeline to extract data from multiple phylogenies is the same as for the single phylogeny example, but now `extract_island_species()` is called for each of the three available phylogenies (i.e. Hawaiian Silversword alliance, *Bidens*, and *Hesperomannia*), with the argument `island tbl` to bind to the same `island_tbl` object.

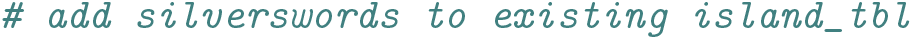
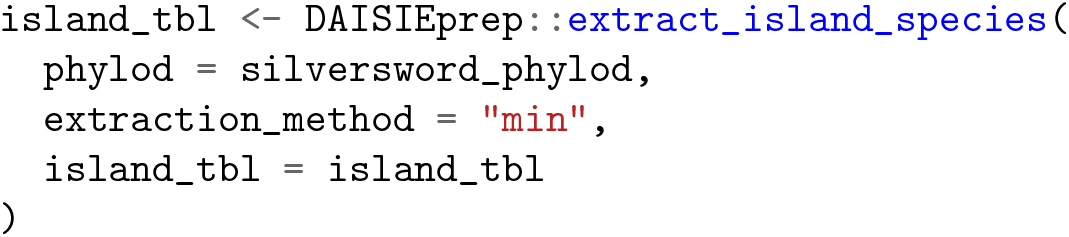

Island colonists that do not have phylogenetic data, but may have a known stem and crown age, are input using the function `add_island_colonist()`. Here the genus *Artemisia* is inserted with a stem and crown age estimate obtained from the literature (Hobbs & Baldwin 2013). The *Lipochaeta-Melanthera* alliance are inserted with a stem age estimated from the literature (Edwards *et al*. 2018) but unknown crown age.

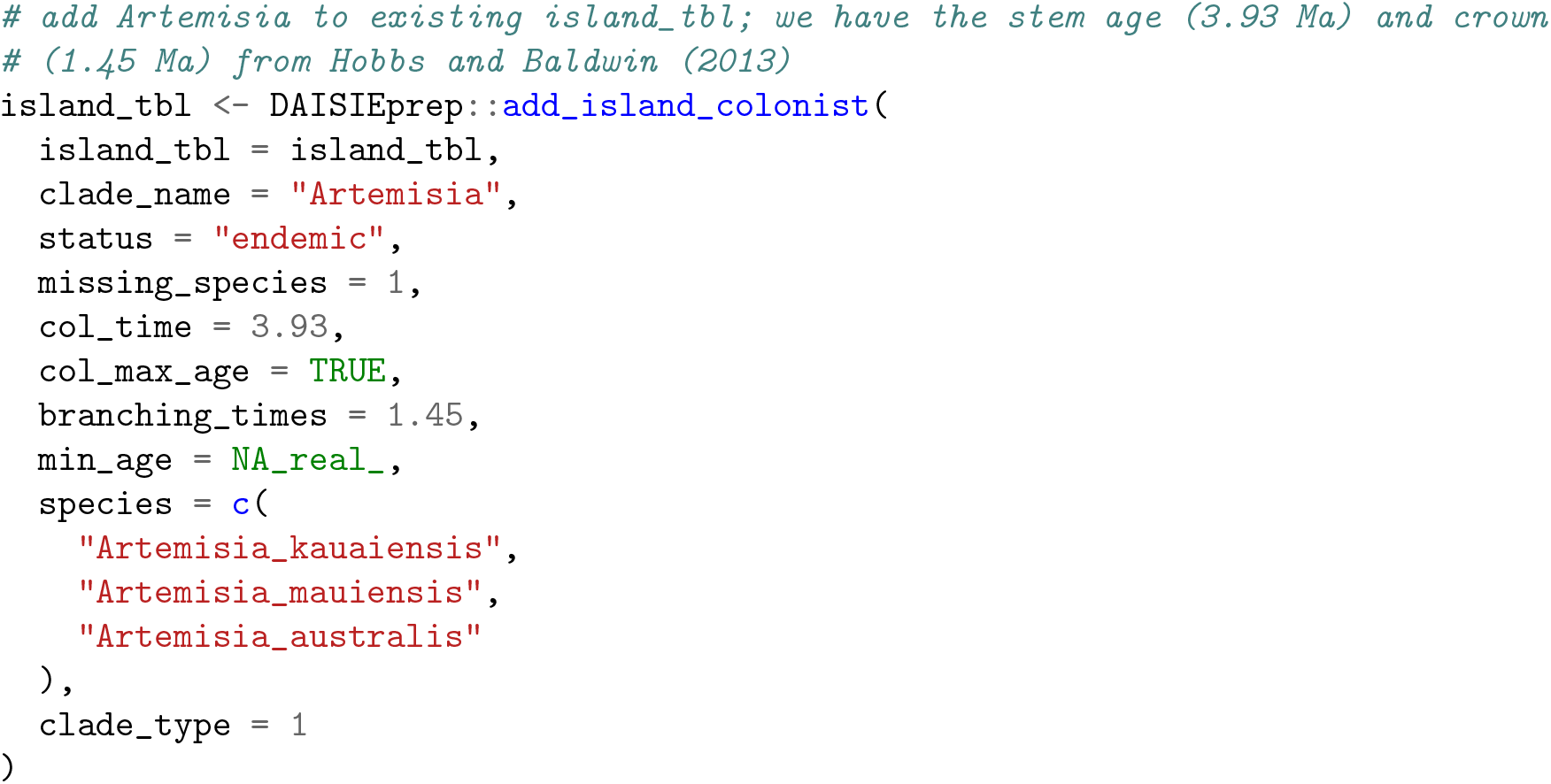

Lastly, when no phylogenetic data is available for a taxonomic group and there exists no estimate for the colonisation time, a clade can be added with the maximum time of colonisation as the age of the island, and the number of species inserted from diversity data (Figure 4). This is done for the *Keysseria* clade in our example.

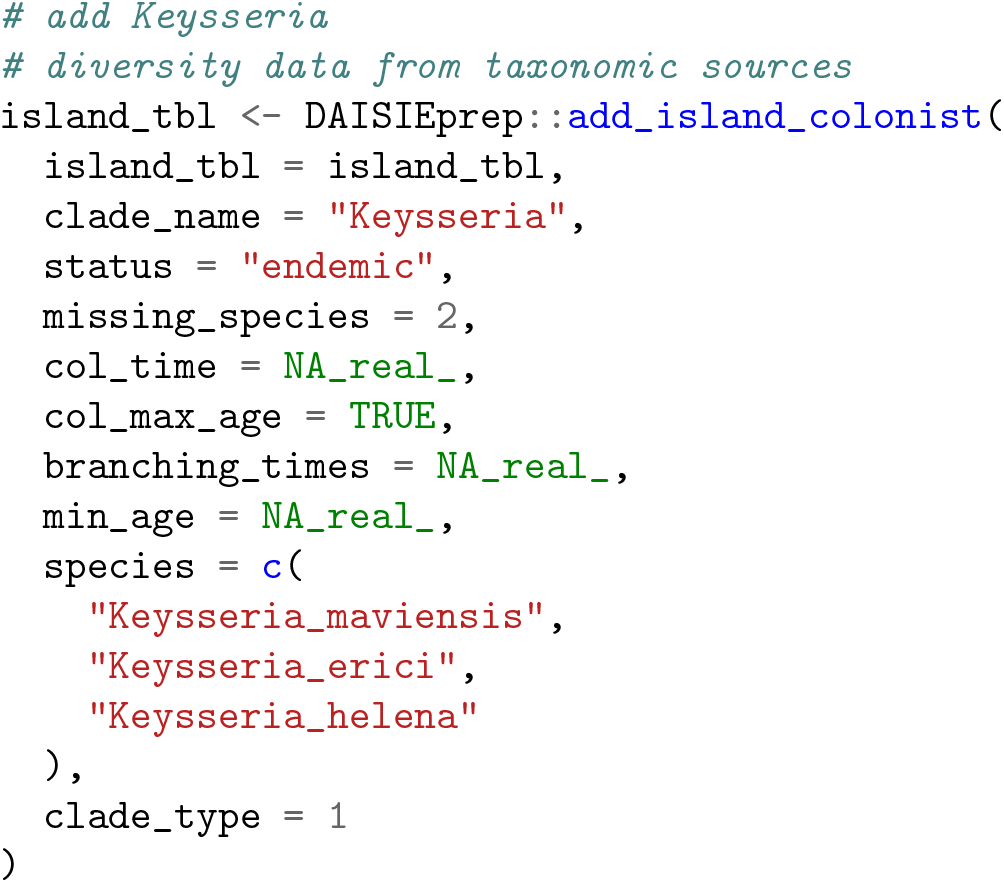

Once a final `Island_tbl` is compiled, `create_daisie_data()` can be used to convert the data for application in DAISIE.

### Posterior distribution example

Phylogenies are estimates of evolutionary history with uncertainty in both branching times and topology, which, in Bayesian analyses, can be represented in a posterior distribution (Huelsenbeck *et al*. 2000; 2001). When available, posterior distributions are often used to repeat downstream analyses on multiple sampled phylogenies, and account for the effect of phylogenetic uncertainty on the conclusions. `DAISIEprep` offers tools to explore the effect of phylogenetic uncertainty on the inferred colonisation and diversification history of the island, by handling the posterior distribution of phylogenetic trees. Here we use the posterior (or ‘pseudo-posterior’, Kuhn *et al*. (2011)) distribution of mammal phylogenies from Upham *et al*. (2019). The island data is individually processed for each phylogeny from the posterior, using `multi_extract_island_species()`, which returns a `Multi_island_tbl` object.

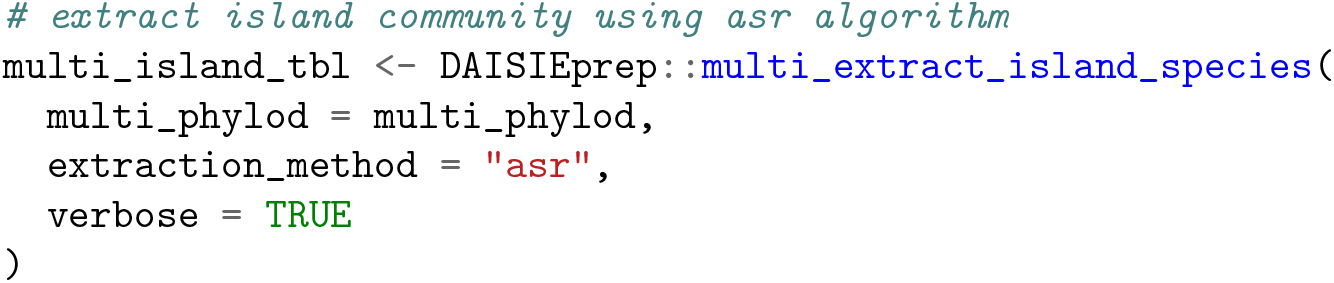

Thus multiple island data sets that reflect phylogenetic uncertainty are produced, with each providing an evolutionary hypothesis for the colonisation and diversification history of the island. Missing species are handled in the same way as for the single phylogeny example that used the maximum clade credibility mammal phylogeny. Finally, `create_daisie_data()` and DAISIE::DAISIE_ML_CS() can be looped over each island data set in order to incorporate the phylogenetic uncertainty. The full script is supplied in the `DAISIEprepExtra` R package.

## Utility functions

The package contains a set of utility functions to inspect the data and check for certain characteristics that may influence the choice of extraction algorithm (‘min’ or ‘asr’). The plotting functions `plot_phylod()` and `plot_colonisation()` are for plotting the data before and after extraction, respectively using the `phylod` object (see above) containing phylogenetic and endemicity data. For inspecting the phylogenetic tree and its associated data, either at the tips, or both at the tips and the internal nodes (Figure 5A), `plot_phylod` provides a way to visualise the distribution of island endemicity across each tree and infer when island colonisation likely occurred. `plot_colonisation` displays for each island colonist lineage their stem age, and if available, the crown age (Figure 5B). `DAISIEprep` uses `ggplot2` (Wickham 2016) and `ggtree` (Yu *et al*. 2017; 2018) to produce plots of the island presence/absence at the tips of the phylogeny, the ancestral states at the nodes and other useful visualisation to aid the user in understanding the data extraction. By building on the flexibility of the `ggplot2` framework `DAISIEprep` plots can be modified using `ggplot2` layers.

**Figure 5:**
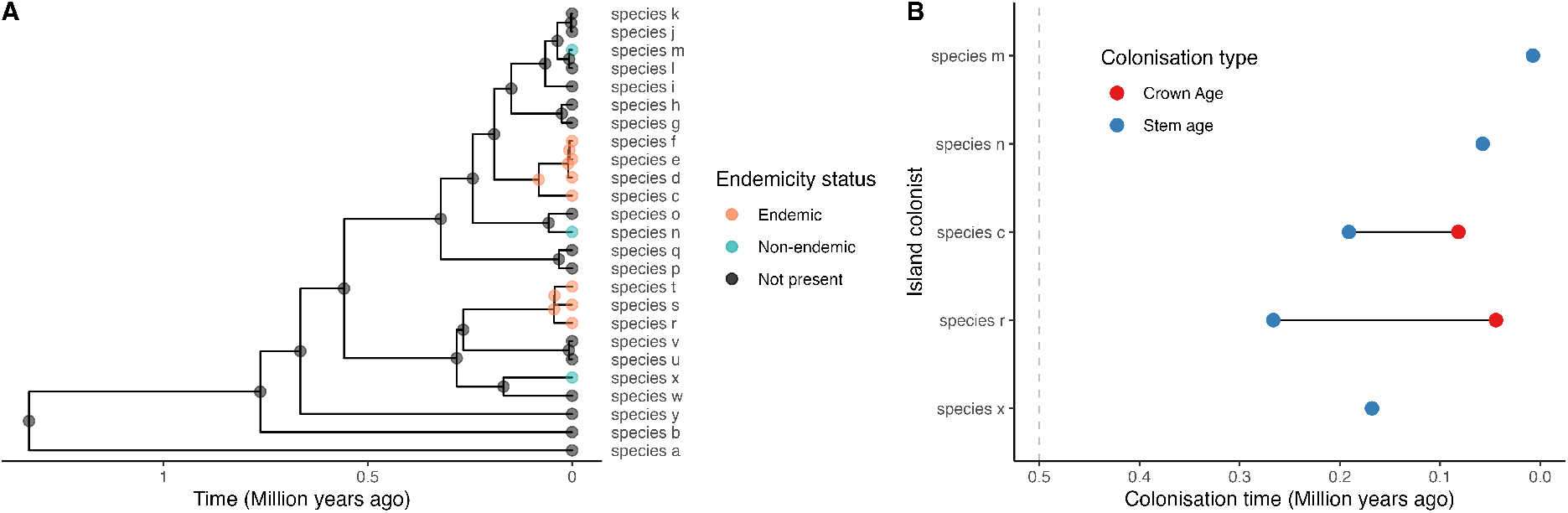
Demonstration of plots produced using (A) `plot_phylo()` and (B) `plot_colonisation()` for the same data. The example shows five island colonists, two of which are island radiations and thus have a crown age as well as a stem age shown in panel B. The dashed line in panel B is the island age.

Another utility function is the ability to check for back-or-onward-colonisation (`any_back_colonisation()`) events within the phylogeny given an ancestral state reconstruction of the island presence. Identifying such colonisations of island species back to the mainland or to another island (thereby changing the endemicity status to non-endemic) can be useful to understand the dispersal patterns of a group and may provide insight into the prevalence of so-called boomerang colonisations (Bellemain & Ricklefs 2008). Identifying these colonisations can be challenging given the inherent limitation of reconstructing species ranges over millions of years, and the specified ancestral state reconstruction model and transition matrix may erroneously produce or miss such colonisations.

Finally, a check for any polyphyly (`any_polyphyly()`) on the tree is implemented in the package, to identify cases in which multiple samples of conspecific species are not found to be monophyletic in the tree. This could for example result from low phylogenetic resolution or incomplete lineage sorting of recently colonised island species, or may indicate true polyphyly (e.g. multiple colonisation events of the island by the same species). When polyphyly of a species is identified, the current extraction algorithms cannot correctly extract these data, so the user needs to decide how to assign colonisation events for that species beforehand.

## Sensitivity of parameter estimates to data extraction method

The objective of using `DAISIEprep` as a processing tool in the pipeline of island biogeography analyses is most often going to be to estimate parameters of colonisation, speciation and extinction dynamics on islands. However, given the different extraction algorithms (‘min’ and ‘asr’) as well as the options within the ‘asr’ algorithm (e.g., model of ancestral range reconstruction and possible geographical transition matrix specified), there are many combinations of extraction settings. These may in some cases produce identical island data sets, while in others they may differ in the number of island colonisation events or branching times. We performed a simple sensitivity analysis (*sensu* global sensitivity, Saltelli *et al*. (2008)) to highlight how data extraction choice can have downstream effects in the parameter estimates obtained by DAISIE. We recommend users to carry out sensitivity analysis tailored to their data, in order to understand how the data extraction method influences outcomes on parameter estimation and model selection. Sensitivity to uncertainty in the island age and the number of mainland species should also be tested. See ‘Sensitivity’ vignette in the `DAISIEprep` package for more details and instructions for running sensitivity analyses.

To illustrate the impact of extraction methods on parameter estimates, we used 100 phylogenies from the posterior of the macro-phylogeny of mammals used in the first example above, focusing again on the species of Madagascar. We extracted the data using the ‘min’ and ‘asr’ methods. For the ‘asr’ method we use two approaches, maximum parsimony and Mk model (assuming equal transition rates) using the R package `castor`. We then applied a diversity-dependent DAISIE model to the resulting datasets, optimising all five parameters. Figure 6 shows that the estimates of anagenesis, cladogenesis and extinction largely overlap, but colonisation rate differs between the methods. This is because the ‘asr’ algorithm produces fewer colonisation events.

**Figure 6:**
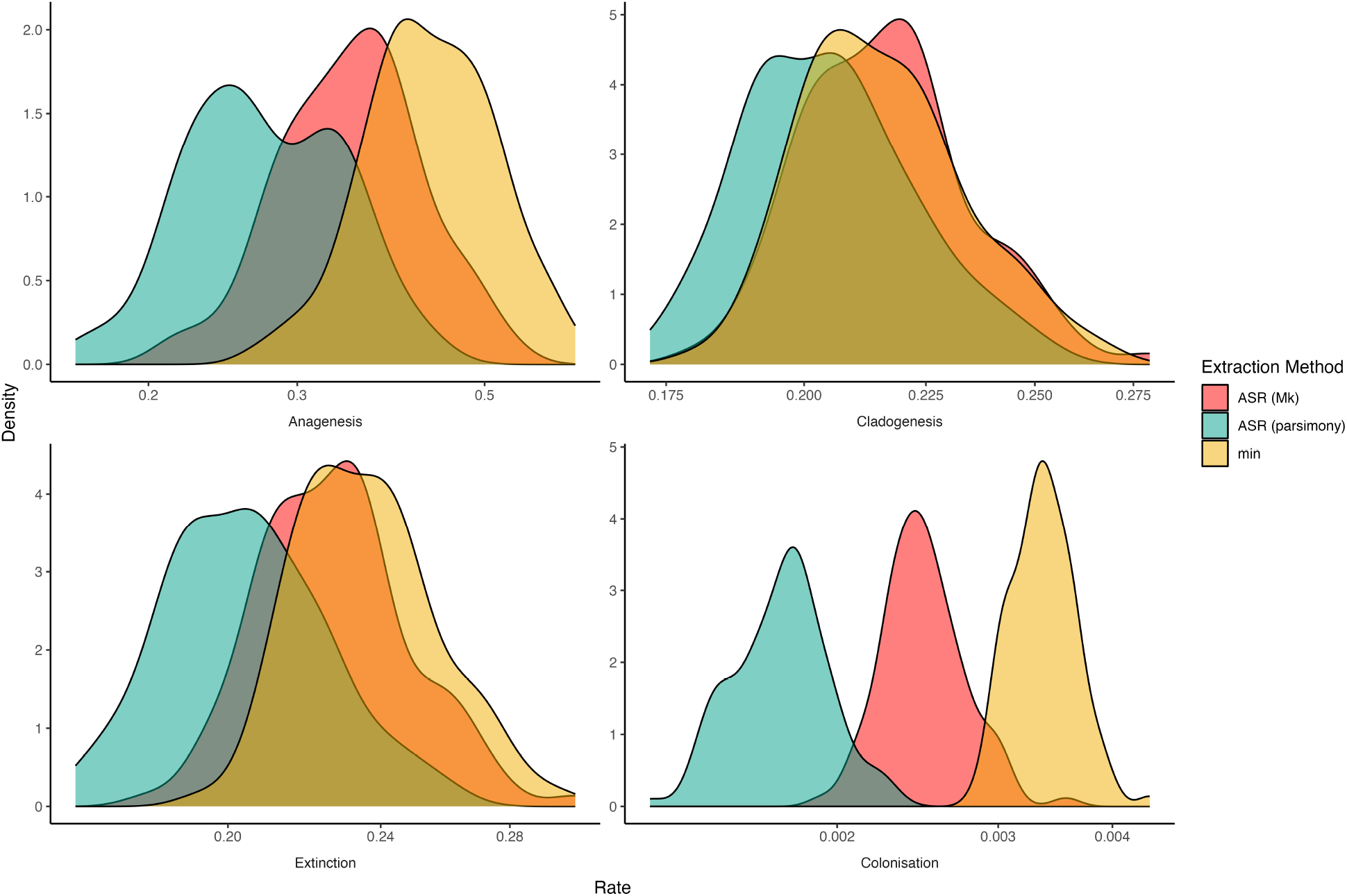
Parameter estimates from the DAISIE maximum likelihood model when data is extracted using different algorithms (‘asr’ and ‘min’) and variants of the ‘asr’ algorithm (parsimony and Mk model). Based on fitting the DAISIE model to 100 datasets (from the posterior distribution of the mammal macro-phylogeny) extracted under each of the extraction algorithms.

## Discussion

Previously, the process of compiling phylogenetic island data to fit community island biogeography models required manually locating island clades. Here we have presented the `DAISIEprep` R package, which automates this procedure and thus allows for a more seamless pipeline to go from commonly used phylogenetic data to model fitting. The examples presented above demonstrate the versatility of the package with a variety of input data that can be handled. The package can be used both on newly inferred phylogenies from sequence data, or previously published phylogenies, for example from a database of phylogenies (e.g. TreeBase, Piel *et al*. (2000)) or macro-phylogenies (Jetz *et al*. 2012; Jetz & Pyron 2018; Upham *et al*. 2019). The performance of the central function (`extract_island_species()`), in terms of computational time is found to be reasonable for common empirical trees (10 - 10,000 tips) with extraction taking seconds. The time consumption of computing the ancestral reconstruction or estimating DAISIE maximum likelihood parameters is on the same time scale or greater, and so this pre-processing tool does not constitute a bottleneck to the pipeline.

Even with this wealth of phylogenetic data there are still many cases for which no phylogenetic data is available or cannot be accessed (Magee *et al*. 2014). In such cases, the absence of a phylogeny can be mitigated by `DAISIEprep`, using island diversity and endemism data in a flexible framework. Using as much phylogenetic data available as possible is evidently still preferred, as it does improve inference reliability (Valente *et al*. 2018), and it improves knowledge on the number of colonisation events.

Both algorithms introduced in `DAISIEprep` (‘min’ and ‘asr’) include ancestral state reconstruction, either implicitly or explicitly. `DAISIEprep` has been set up to allow for alternative methods of ancestral state reconstruction methods. Therefore, if it is perceived that species range, either on the island, on the mainland or both, influences the rates of speciation and extinction, then a model that can account for these causes of rate heterogeneity can be used (Maddison 2006; Maddison *et al*. 2007; Holland *et al*. 2020). However, it is worth noting the known limitations of these methods. Particularly, they are highly uncertain further back in time (Cunningham 1999; Martins 2000; O’Meara & Beaulieu 2021). One encouraging aspect is that islands – especially oceanic islands – are usually young (5-10 Mya) and so species will have colonised relatively recently, making the inference of species arrival more accurate than if it were deeper in the past. Another limitation is the use of phylogenies containing species for which no molecular data is available and which are added to the trees using taxonomic constraints. Such methods have been shown to yield reliable, albeit conservative, conclusions from diversification analyses (Chang *et al*. 2020), yet, these methods may scrabble the phylogenetic signal of island presence, especially when these missing species are only constrained at higher taxonomic levels of family or order (Rabosky 2015). In turn, this may potentially lead to an increase in island colonisation events as species from the same insular radiation are incorrectly placed in different locations across the tree. In these cases, results should be validated with the equivalent DNA-only phylogeny.

By releasing this open-source package we hope to encourage the use, development and scrutiny of phylogenetic inference models, and allow the community of phylogenetic inference to expand and answer more questions in the fields of biogeography and community assembly, while providing high reproducibility. This package will continue development in parallel with the `DAISIE` R package.

## Acknowledgments

We would like to thank the Theoretical and Evolutionary Community Ecology group at the University of Groningen, particularly Pedro Santos Neves for discussions and their input. We would like to thank the Center for Information Technology of the University of Groningen for their support and for providing access to the Peregrine high performance computing cluster. JWL was funded through a Study Abroad Studentship by the Leverhulme Trust and was also funded by a NWO VICI grant awarded to RSE who was funded by the same grant. TP was funded by the University of Groningen and the Faculty of Natural Sciences, University of Stirling (UK). LR was funded by an NWO VIDI grant awarded to LV, LV was funded by the same grant. All authors declare no conflict of interest.

## Authors contributions

JWL formulated the project, drafted the manuscript, created and led development on the `DAISIEprep` package, and ran the analyses. LR collected the Hawaiian Asteraceae data and assisted in writing the R script and manuscript on the multiple phylogeny example. TP wrote the extending ancestral state reconstruction vignette in the R package. LV collected the data for the Malagasy Mammals (originally from Michielsen *et al*. (2023)) and assisted in the development of the R package, as well as supervised the project. RSE supervised the project and gave feedback throughout. All authors revised the manuscript and approved publication.

